# PICKLUSTER: A protein-interface clustering and analysis plug-in for UCSF ChimeraX

**DOI:** 10.1101/2023.06.19.545106

**Authors:** Luca R. Genz, Thomas Mulvaney, Sanjana Nair, Maya Topf

**Affiliations:** Leibniz-Institut für Virologie (LIV), Hamburg, Germany; Universität Hamburg, Hamburg, Germany; Universitätsklinikum Hamburg Eppendorf (UKE), Hamburg, Germany; Centre for Structural Systems Biology (CSSB), Hamburg, Germany

## Abstract

**Motivation:** The identification and characterization of interfaces in protein complexes is crucial for understanding the mechanisms of molecular recognition. These interfaces are also attractive targets for protein inhibition. However, targeting protein interfaces can be challenging for large interfaces that consist of multiple interacting regions. We present PICKLUSTER -a program for identifying sub-interfaces in protein-protein complexes using distance clustering. The division of the interface into smaller “sub-interfaces” offers a more focused approach for targeting protein-protein interfaces.

**Availability and implementation:** The plug-in PICKLUSTER is implemented for the molecular visualization program UCSF ChimeraX 1.4 and subsequent versions and and is freely available in the ChimeraX toolshed or, together with the source code, from https://gitlab.com/topf-lab/pickluster.git).

## Introduction

Protein complexes are key components of the majority of biological processes within cells such as the activation of receptor molecules, DNA replication, signal transduction, protein export and transport, recombination and repair (Spirin and Mirny, 2003; Jubb *et al*., 2015; Yan and Wang, 2012). Understanding the mechanisms of protein complex formation and macromolecular recognition requires the identification and characterization of protein-protein interfaces (Xue *et al*., 2015). Additionally, these protein interfaces can be targeted for protein complex inhibition, which is an approach in drug discovery (Jubb *et al*., 2015; Oltersdorf *et al*., 2005).

Protein interfaces are typically identified through high-resolution structures of protein complexes that are generated using experimental methods such as X-ray crystallography or cryo-electron microscopy (cryo-EM) (Xue *et al*., 2015). Due to the inherent nature of these experiments, large-scale structure determination is not a viable option. Recent advances in AI-based structure prediction methods, notably AlphaFold2 (Jumper *et al*., 2021), help in circumventing this problem by providing the possibility for large scale structure prediction with accuracy often close to what is achieved by experiments.

The analysis of protein complexes generated by either experiments or predictions has revealed that protein-protein interfaces can be of different sizes (Yan *et al*., 2008). Large protein interfaces can consist of multiple interacting domains that are geometrically separated, posing a challenge for targeting the entire interface using drugs (Blundell *et al*., 2000, 2006). Moreover, previous research has shown the importance of small binding pockets in the protein interface to increase the selectivity of protein interface binders (Blundell *et al*., 2006; Jubb *et al*., 2012). The division of the interface into smaller sub-interfaces based on their spatial properties could facilitate the targeting of these interfaces.

In this study, we have developed PICKLUSTER (Protein Interface C(K)luster), a UCSF ChimeraX-1.4 (Pettersen *et al*., 2021) plug-in that clusters protein interfaces based on distance. PICKLUSTER provides various scoring metrics for the analysis of the interface, not only of structures of protein complexes but also of models generated by AlphaFold2. By fragmentation of the protein interface, it offers a focused and useful approach for targeting protein-protein interfaces.

## Implementation

PICKLUSTER is written in python 3.9 and implemented as a UCSF ChimeraX 1.4 plug-in. To calculate the interface, it requires as input the PDB/mmCIF file of a protein complex, the chain identifiers for the pair of chains and the model classification (*Experimental protein structure/AlphaFold2 model*). In case of an AlphaFold model, the *JSON* or *Pickle* file is also required as an input. PICKLUSTER colors the protein complex according to protein interface clusters in the input chains and reports the list of the residues at the interface in the UCSF ChimeraX Log.

### Clustering algorithm

For the efficient identification of interfaces, the distances between heavy atoms (non-hydrogen atoms) user-specified chains are calculated using a KDTree data structure (provided by the *SciPy* python package (Virtanen *et al*., 2020)). Heavy atoms within a distance of 5.0Å between the chains are considered “interaction partners” (Evans *et al*., 2022). For AlphaFold2 models, the residues exhibiting a predicted local distance difference test (pLDDT) (Jumper *et al*., 2021; Akdel *et al*., 2022) score below the cut-off (default: 50) are excluded from the interface. Subsequently, the center of mass of each residue exhibiting an “interaction partner” in the protein interface is computed. The clustering of the interface is performed by calculating the distance of these centers of masses in the interface. If the distance is below the distance cut-off (default: 5.0Å) the residues are considered to be in the same cluster.

### Features of PICKLUSTER

To illustrate the capabilities of PICKLUSTER, we employed it to identify the interface of a structural model generated by the ColabFold tool, which is based on AlphaFold2 (Mirdita *et al*., 2022). We focused on the interface of UL15 and UL33 in a heterotrimeric complex with UL28 (for full complex see Fig. S1) from *Epstein-Barr virus* (EBV). The output provides various types of structural information, including the detection of three clusters (Fig. 1A). The plug-in offers the option of visualizing the clusters in the protein sequence as it generates a Sequence Coloring Format (SCF) file that can be imported into the UCSF ChimeraX Sequence viewer (Fig. 1B). For the analysis of protein interfaces in models generated by AlphaFold2, the plug-in provides the option to display the clusters colored by their pLDDT (Fig. 1C), a predicted aligned error (PAE) matrix (Jumper *et al*., 2021), the max. PAE of the model, the mean PAE per cluster, the predicted template modeling (pTM) score and the interface pTM (ipTM). To access these scores, PICKLUSTER requires a file in JSON or Pickle format that is obtained during the structure prediction. Regardless of the input structure-type category (AlphaFold2 or Experimental Structure), PICKLUSTER also offers the option to display a list of interacting residues per cluster in the UCSF ChimeraX Log and can show the atomic interactions in the protein complex visualization in the UCSF ChimeraX main window (Fig. 1D) (complete list of options is shown in Fig. 1E).

**Fig. 1.**
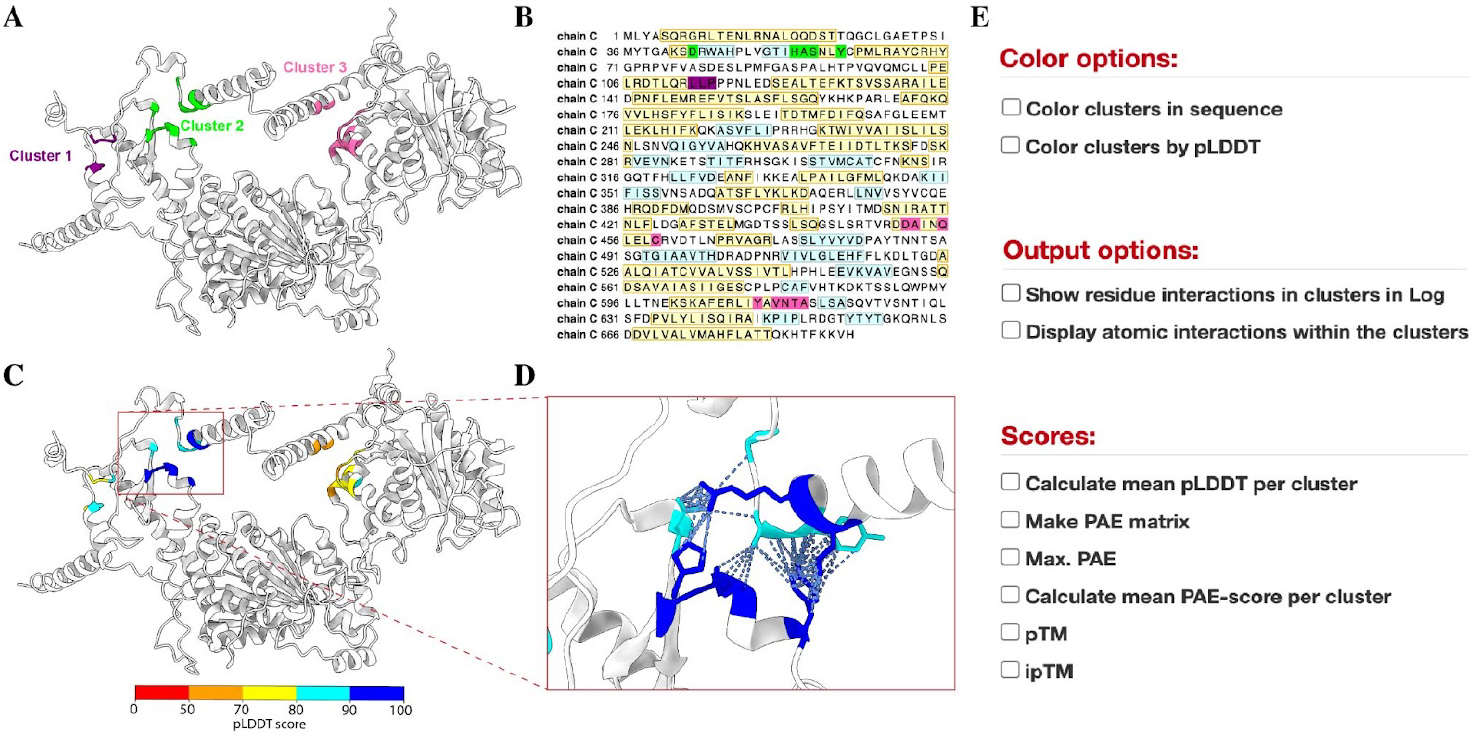
Representative example of PICKLUSTER outputs applied to the dimeric complex of UL15 and UL33 from Epstein-Barr Virus. **(A)** PICKLUSTER output for UL15 and UL33 in a trimeric complex with UL28 (full complex is shown in Fig. S1) from Eppstein-Barr virus (EBV) generated using Colabfold with the three protein interface residue clusters colored red,green and blue. The clusters are calculated using a distance cut-off of 5Å for the clustering. **(B)** Sequence of chain C with the residues colored according to the clusters. **(C)** The clusters in (A) are colored according to their pLDDT with red indicating regions of poor confidence and blue depicting high confidence regions. **(D)** Inset of (C) displaying the atomic interactions in cluster 2 (as in (A)). **(E)** Options provided by PICKLUSTER for the visualization and analysis of the clustered protein interface.

In this example of UL15 and UL33 from EBV, we observed that the pLDDT at the interface cluster 2 (Fig. 1D) is high and the error is low (mean PAE: 2.26Å), which indicates high confidence in the predicted interface cluster.

## Conclusion

We have developed a new plug-in for ChimeraX – PICKLUSTER, which clusters protein interfaces based on spatial properties and provides a range of scoring metrics for the analysis of these interfaces. The smaller clusters predicted by the program can be useful for many applications, such as ligand docking, analysis of hot-spot residues, functional characterization of mutations and understanding the mechanisms of protein association. Moreover, by facilitating the identification of the interface, PICKLUSTER can potentially be a useful tool in protein modeling challenges like CASP (Critical Assessment of protein Structure Prediction) (Kryshtafovych *et al*., 2019) if the focus is on protein interfaces. Additionally, mapping the sub interfaces onto the sequence of protein complexes holds the potential to discover protein complexes with a similar interaction pattern.

## Acknowledgements

The authors thank the Topf group for discussions and valuable feedback.

## References

Akdel, M. et al. (2022) A structural biology community assessment of AlphaFold2 applications. Nat. Struct. Mol. Biol., 29, 1056–1067.

Blundell, T.L. et al. (2000) Protein-protein interactions in receptor activation and intracellular signalling. Biol. Chem., 381, 955–959.

Blundell, T.L. et al. (2006) Structural biology and bioinformatics in drug design: opportunities and challenges for target identification and lead discovery. Philos. Trans. R. Soc. Lond. B Biol. Sci., 361, 413–423.

Evans, R. et al. (2022) Protein complex prediction with AlphaFold-Multimer. bioRxiv, 2021.10.04.463034.

Jubb, H. et al. (2015) Flexibility and small pockets at protein–protein interfaces: New insights into druggability. Prog. Biophys. Mol. Biol., 119, 2–9.

Jubb, H. et al. (2012) Structural biology and drug discovery for protein–protein interactions. Trends Pharmacol. Sci., 33, 241–248.

Jumper, J. et al. (2021) Highly accurate protein structure prediction with AlphaFold. Nature, 596, 583–589.

Kryshtafovych, A. et al. (2019) Critical assessment of methods of protein structure prediction (CASP)-Round XIII. Proteins, 87, 1011–1020.

Mirdita, M. et al. (2022) ColabFold: making protein folding accessible to all. Nat. Methods, 19, 679–682.

Oltersdorf, T. et al. (2005) An inhibitor of Bcl-2 family proteins induces regression of solid tumours. Nature, 435, 677–681.

Pettersen, E.F. et al. (2021) UCSF ChimeraX: Structure visualization for researchers, educators, and developers. Protein Sci., 30, 70–82.

Spirin, V. and Mirny, L.A. (2003) Protein complexes and functional modules in molecular networks. Proc. Natl. Acad. Sci. U. S. A., 100, 12123–12128.

Virtanen, P. et al. (2020) SciPy 1.0: fundamental algorithms for scientific computing in Python. Nat. Methods, 17, 261–272.

Xue, L.C. et al. (2015) Computational prediction of protein interfaces: A review of data driven methods. FEBS Lett., 589, 3516–3526.

Yan, C. et al. (2008) Characterization of Protein–Protein Interfaces. Protein J., 27, 59–70.

Yan, Z. and Wang, J. (2012) Specificity quantification of biomolecular recognition and its implication for drug discovery. Sci. Rep., 2, 309.

