## Supplementary Materials for "PICKLUSTER: A protein-interface clustering and analysis plug-in for UCSF ChimeraX"

**The PDF file includes:**

Fig. S1

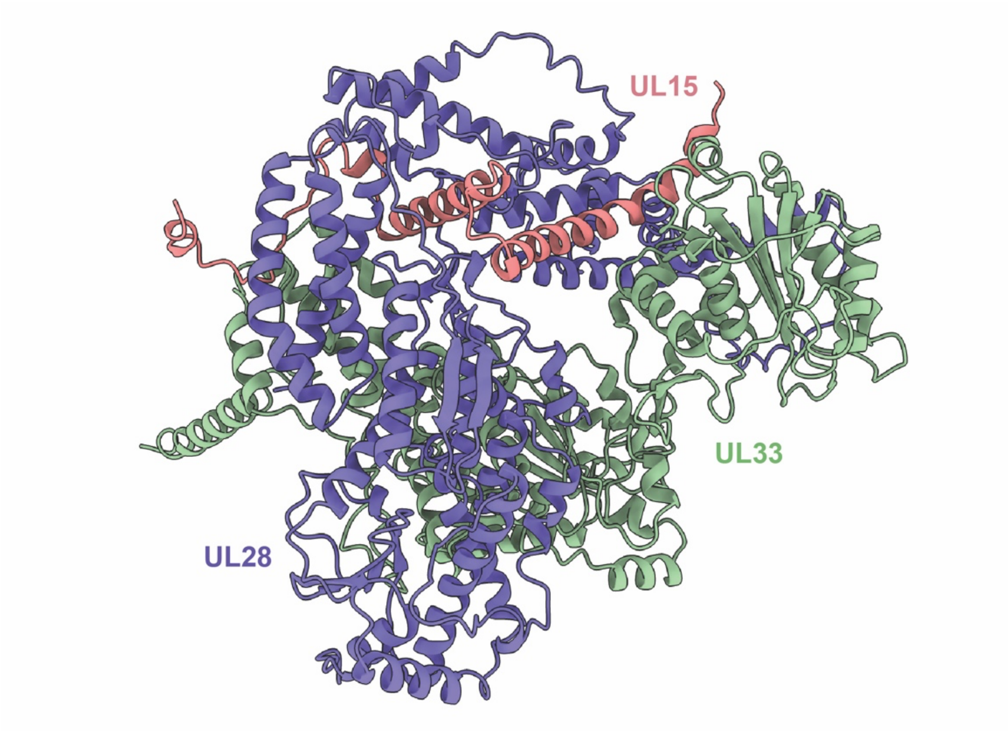


**Fig. S1.** Trimeric complex of UL28 (purple), UL15 (salmon) and UL33 (green) from *Epstein-Barr Virus* modelled with ColabFold (Mirdita *et al.*, 2022).
